# >Corna - An Open Source Python Tool For Natural Abundance Correction In Isotope Tracer Experiments

**DOI:** 10.1101/2020.09.19.304741

**Authors:** Raaisa Raaisa, Shefali Lathwal, Victor Chubukov, Richard G. Kibbey, Abhishek K. Jha

## Abstract

**Background:** Stable isotope-based approaches are used in the field of metabolomics for quantification and identification of metabolites, discovery of new pathways and measurement of intracellular fluxes. In these experiments, often performed with mass spectrometry (MS), data must be corrected for natural abundance of isotopes. Various stand-alone tools with their own separate data formats and learning curves exist for correction of data collected at different resolutions, for tandem MS, and for different number of tracer elements.

**Results:** We present a Python package, Corna, that combines natural abundance correction workflows for several experimental conditions and can be used as a one-stop-shop for stable isotope labeled experiments. We validate the algorithms in Corna with published tools, where available, and include new features, such as correction of two tracer elements, that are not yet implemented in any existing software application as per our knowledge. We also present the integration of Corna with an existing open source peak integration software. The integrated workflow can reduce processing times for a typical stable isotope based workflow from days to hours for a familiar user.

**Conclusions:** Algorithmic advancements have been keeping up with the developments in mass spectrometry technologies and have been the focus of most existing tools for natural abundance correction. However, in this high throughput era, it is also important to recognize user experience, and integrated and reproducible workflows. Corna has been written in Python and is designed for users who have access to large amounts of data from different kinds of experiments and want to integrate a natural abundance correction tool seamlessly in their pipelines. The latest version of Corna can be accessed at https://github.com/raaisakuk/NA_Correction.

## Introduction

Stable isotope based approaches are widely used in metabolomics to study dynamic changes within the metabolome, quantify and identify metabolites, discover new pathways, and measure metabolic fluxes in cells [1]. These approaches often use mass spectrometry (MS) coupled with liquid chromatography (LC) or gas chromatography (GC) to track the flow of isotopes of common elements such as C, H, and N (referred to as tracer elements) from a substrate through a metabolic network.

A fraction of most elements used in isotope tracer studies occur naturally in their isotopic forms. Therefore, the analysis of data from a stable isotope tracer experiment should remove the contribution of the naturally occurring isotopes. This correction is commonly referred to as the natural abundance correction. Natural abundance correction is especially crucial in quantitative approaches like metabolic flux analysis where calculations of absolute fluxes through metabolic enzymes are highly sensitive to the isotopic enrichment values. [2].

Brauman [3] implemented a method for natural abundance correction of MS data in the programming language ALGOL and recommended a software [4] that could be used for such computations as far back as 1966. Using probability and combinatorics, various algorithms have been developed for correcting natural abundance from MS [5]–[9] and MS/MS [10]–[12] data. Several software packages have been written to implement these algorithms. One of the most widely used tools, IsoCor [13], [14] is written in Python and has a graphical user interface (GUI). It can process MS data and can correct for naturally occurring isotopes of elements other than the tracer element, at different resolutions. FluxFix [15] is another platform for correcting MS data that provides a GUI. This web-based platform can correct single isotope labeled experiments and uses experimental data from unlabeled samples for natural abundance correction. However, both FluxFix and IsoCor cannot process MS/MS data or data containing multiple tracer elements simultaneously. Another Python-based software, PyNAC, can perform correction with multiple isotope tracer elements [16], but cannot handle tandem MS or low resolution MS data. AccuCor [17], written in R, can process MS data at different resolutions, but cannot process MS/MS data or handle multiple tracer elements. For tandem mass spectrometry data, there are software packages such as Isotope Correction Toolbox written in Perl [12] and PIDC [10] written in MATLAB. Unlike applications written in Python, R, or Perl, which are open-source programming languages, applications written in MATLAB require a license that can be expensive and, therefore, infeasible to obtain for everyone. Additionally, none of the above tools track metadata associated with the samples and metabolites in the experiment during processing, which is a critical gap since these metadata are crucial for downstream analyses and visualizations.

In this paper we introduce a Python package, “Corna”, with the goal of reducing the need to employ different tools for natural abundance correction. We implement algorithms for LCMS/GCMS/LCMS-MS data at various resolutions with focus on a modular structure that allows easy integration with other tools in stable isotope labeled metabolomics data processing pipelines. A comparison of Corna with other natural abundance correction packages and software tools is given in Table 1.

**Table 1:**
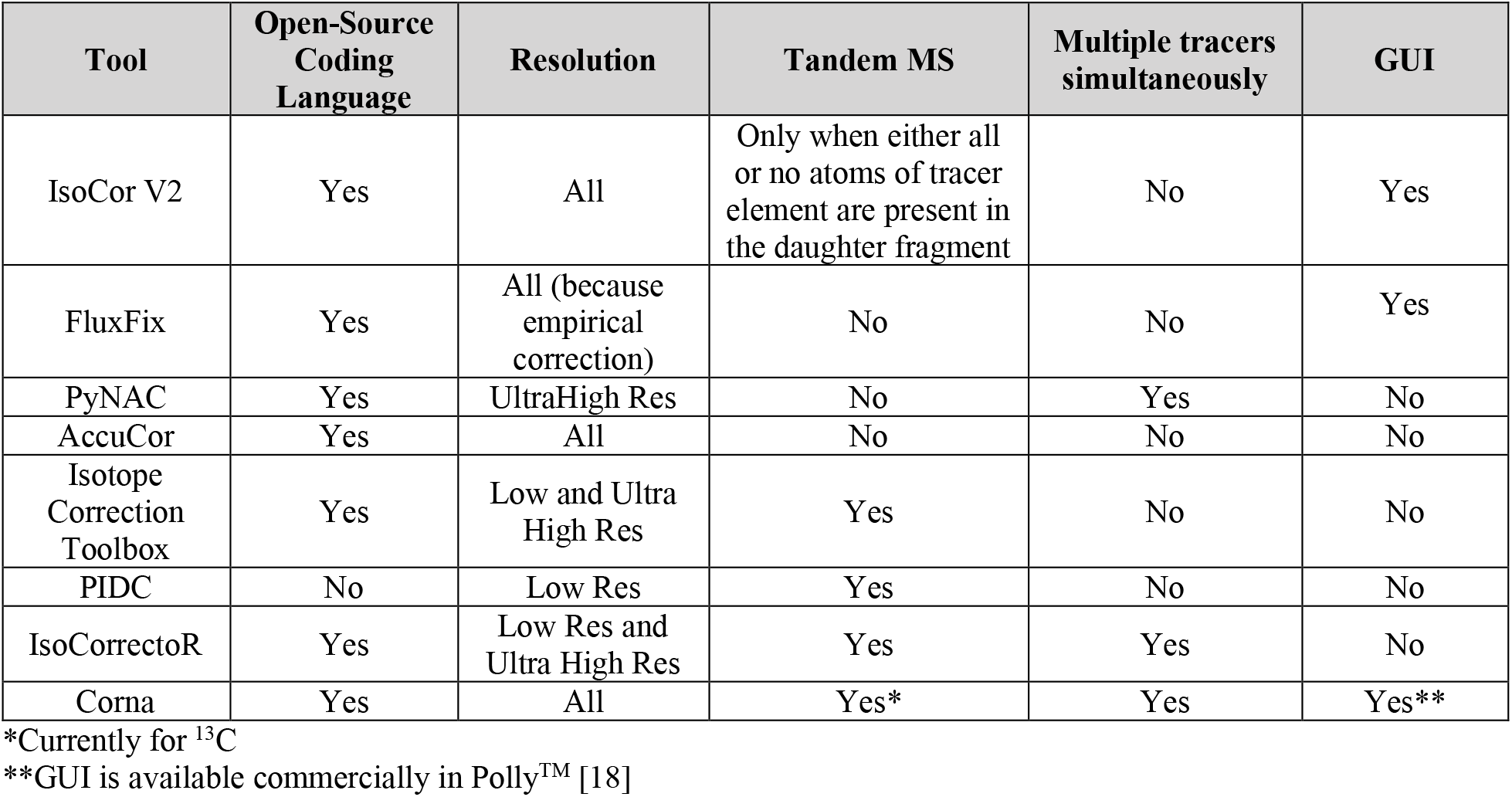
Comparison of features in Corna and available natural abundance correction tools

| Tool | Open-Source Coding Language | Resolution | Tandem MS | Multiple tracers simultaneously | GUI |
| --- | --- | --- | --- | --- | --- |
| IsoCor V2 | Yes | All | Only when either all or no atoms of tracer element are present in the daughter fragment | No | Yes |
| FluxFix | Yes | All (because empirical correction) | No | No | Yes |
| PyNAC | Yes | UltraHigh Res | No | Yes | No |
| AccuCor | Yes | All | No | No | No |
| Isotope Correction Toolbox | Yes | Low and Ultra High Res | Yes | No | No |
| PIDC | No | Low Res | Yes | No | No |
| IsoCorrectoR | Yes | Low Res and Ultra High Res | Yes | Yes | No |
| Corna | Yes | All | Yes* | Yes | Yes** |

\*Currently for $^{13}\text{C}$
\*\*GUI is available commercially in Polly™ [18]

## Key Capabilities

### Open-source programming language

Corna has been written in an open-source programming language, Python. Python can be run without a license on Mac, Windows and Linux operating systems with equal ease. Therefore, Corna benefits from platform independence. The implementation of the package follows a class-based architecture with an application programming interface (API) that can be used as-is, or wrapped in a user friendly script or web interface. Sample Interactive-python (Ipython) notebooks have been made available with the package (Table 2). These notebooks cover all algorithms and features available in Corna. A web implementation of Corna is also available in Polly™ [18].

**Table 2.** Sample workflows implemented in Ipython notebooks

| Experiment | Label | Resolution | Ipython Notebook |
| --- | --- | --- | --- |
| LCMS/MS | C13 | Ultra High | N1, N8 |
| LCMS | C13 and N15 | Ultra High | N2 |
| LCMS | C13 | Low | N3, N7 |
| GCMS | C13 | Low | N4 |
| LCMS | H2 and N15 | User provided | N5 |
| LCMS | N15 | User provided | N6 |

### Support for different MS workflows and user-friendly features

Corna can be used for data generated from LCMS, LCMS-MS and GCMS experiments. For LCMS and GCMS, it can handle experiments that generate data with single or dual labeled metabolites. In an isotopic tracer experiment, resolution of the mass spectrometer governs whether the masses of naturally occurring isotopes of elements other than the tracer element can be resolved from the tracer. This ability to find indistinguishable isotopes (described in Results section) and include them in the correction is also handled for LCMS and GCMS data. For very low resolution or ultra high resolution mass spectrometer the input can be the terms ‘low res’ and ‘ultra high res’ respectively. For Orbitrap and FT-ICR, resolution and the mass at which that resolution is specified can be entered for auto-detection of indistinguishable isotopes. For LCMS-MS studies, additional background noise can be removed using unlabeled samples, referred to as the background samples in Corna. Once the measured intensities from the experiment have been corrected, Corna calculates fractions for different isotopologues (for LCMS or GCMS) or fragments (for LCMS-MS) and pool totals of metabolites in all samples. Brief descriptions of the algorithms are given in the sections below with details included in the supplementary information (S1).

#### MS

MS algorithms in Corna can be run for experiments with single or dual-labeled metabolites. For correcting single labeled LCMS data, we generate a correction matrix for each metabolite by calculating mass isotopomer distributions of the labeled tracer and its corresponding indistinguishable isotopes from their known natural abundances. We then multiply inverse of the correction matrix with observed intensities [7] as shown in Equation 1.

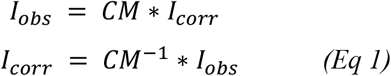

For ultra high resolution data, this algorithm can be extended to dual-labeled metabolites. When there are two distinguishable isotopic tracer elements in the data, Corna calculates independent correction matrices for each isotope and sequentially corrects data corresponding to each tracer. Apart from the tracers, natural abundance of isotopes of elements which are not tracers but present in the compound have to be taken into account. These isotopes, although resolved, are not measured and lead to loss of experimentally-relevant tracer to natural abundance of the unmeasured isotopes, [17]. This correction is also accounted for during in Corna.

For high resolution machines, which can resolve certain isotopes from the tracer element but not all, the correction matrix is populated only with terms that are not resolved [17]. The ability to resolve a naturally abundant isotope from an experimentally labeled tracer element varies with the mass of the metabolite, and the mass difference between the isotope and the element at a given machine resolution. This relationship can be different for different mass spectrometers and has been implemented for Orbitrap [17] and FT-ICR [19].

#### MS with derivatization

In some experiments (e.g. most GC-MS methods), metabolites are derivatized through chemical modifications before measurement. In these experiments, there is a metabolite of interest and a derivatizing agent, which is the chemical modification added on to the metabolite. The atoms of the derivatizing agent are not labeled experimentally but those of the metabolite can be labeled from the experiment. Therefore, natural abundance correction of these data needs to account for two additional factors, i) the mass of the measured molecule consists of both the metabolite and the derivatizing agent, and ii) the metabolite and derivatizing agent can have the same elements in them, but only the metabolite can have isotopically labeled elements from the tracer used in the experiment. Since elements of the derivatizing agent can only be labeled due to natural abundance, Corna treats the derivatizing agent’s atoms like indistinguishable isotopes (Supplementary S1.1.4). Therefore, the algorithm for low resolution LCMS with indistinguishable isotopes can be used to correct data from experiments performed with a derivatizing agent.

#### MS/MS

In a labeled tandem mass spectrometry experiment, mass isotopomer distribution in the MS/MS data is influenced by the labeling pattern in the parent molecules and the elements present in the daughter fragments. This distribution of labels during fragmentation is accounted for C13 tracer experiments by using the formula in equation 2 [20].

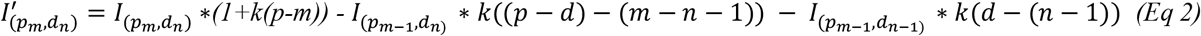

where *m* − *n* ≤ *p* − *d*, *p* and *d* are the total number of carbons in parent ion and daughter ions, respectively, *m* and *n* are number of C13 in parent and daughter ion, *I*(*p*_*m*_,*d*_*n*_) is the peak area corresponding to a daughter ion with mass *n* and parent ion with mass *m*, and *k* is the natural abundance of C13.

### Ease of I/O and Modular architecture

Within Corna, the input/output layer is independent of the processing layer. Simplified architecture of Corna is shown in Figure 1. The architecture consists of four layers; I/O, ‘Parser’, ‘Model’, and ‘Algorithm’. The I/O layer requires information such as the tracer isotope used in the study, chemical formula of each species to be corrected, formula of daughter fragments corresponding to each parent in MS/MS, resolution of the mass spectrometer defined at a given mass, natural abundances of various isotopes and intensity values to be corrected. This information is extracted by the Parser, stored as pythonic data structure by Model, and passed to the Algorithm for processing. The Algorithm layer gives the natural abundance corrected intensities back to the Model. The corrected values are returned to Parser layer and put into a format suitable for display. The separation of layers allows easy adaptability of Corna to different software packages with different output formats. In these cases, only the Parser layer can be modified without affecting the rest of the code. Similarly, different algorithms can be added to the Algorithm layer over time. Separating the I/O layer from the rest of the codebase also helps in adding validation layers to check the input data for formatting and other errors. Corna allows .txt/.csv/.xlsx/.xls formats as input and generates output in .csv format. The .csv file output can be visualized using python libraries such as matplotlib, or programs such as TIBCO Spotfire or Microsoft Excel. Any sample metadata can also be provided by the user along with the data and in these cases, the final output will contain the metadata in the same file as the corrected data. With the current Parser, Corna can directly use the peak table outputs from MultiQuant™ (SCIEX) and El-MAVEN.

**Figure 1.**
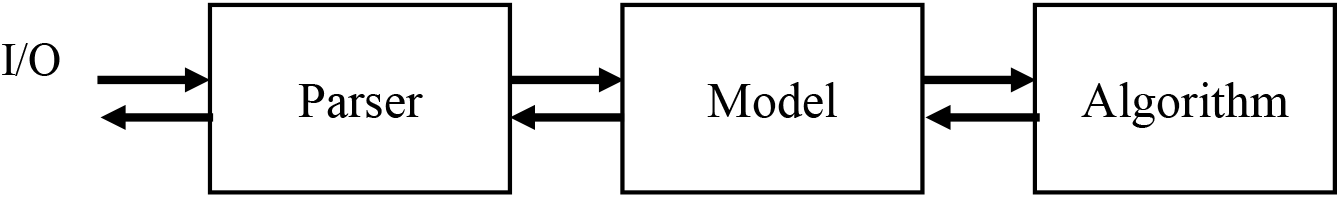
Modular architecture in Corna

## Results

To validate the algorithms implemented in Corna, we compared the output from Corna with published software packages. For single isotope labeled low resolution MS data, we used matrix-based natural abundance correction algorithm [7] and compared the results with IsoCor V1 [13] (Figures 2 and 3). We used the data available in the file “Example single meas.txt” from Examples folder in IsoCor V1 and corrected for C13 tracer assuming that they represent a four-carbon metabolite, succinate. IsoCor V1 considers all isotopes as indistinguishable by default, and we assumed 100% purity of label. The same settings were used in Corna. N15 tracer data were obtained from the data file, N_Sample_Input.xlsx, provided by the package, AccuCor [17]. Figure 2 shows the raw and corrected fractional enrichments (FE) for AcetylCoA from sample N15_10_140k_A using both IsoCor V1 and Corna. For correction of both C13 and N15 tracers, the corrected fractional enrichments from Corna were comparable with IsoCor V1 despite differences in the numerical solvers used for equation (1).

**Figure 2.**
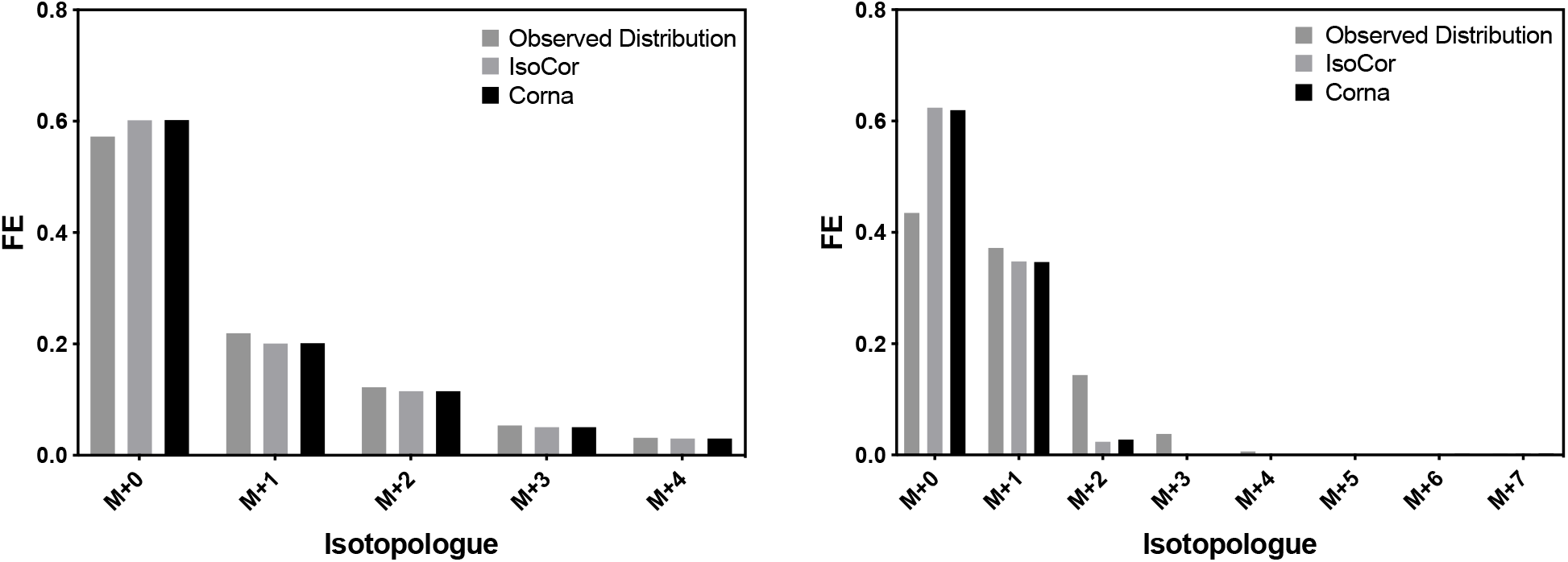
Observed distribution for (a) Succinate labeled with isotope C13 and (b) Acetyl CoA labeled with N15 corrected by IsoCorV1 and Corna

**Figure 3.**
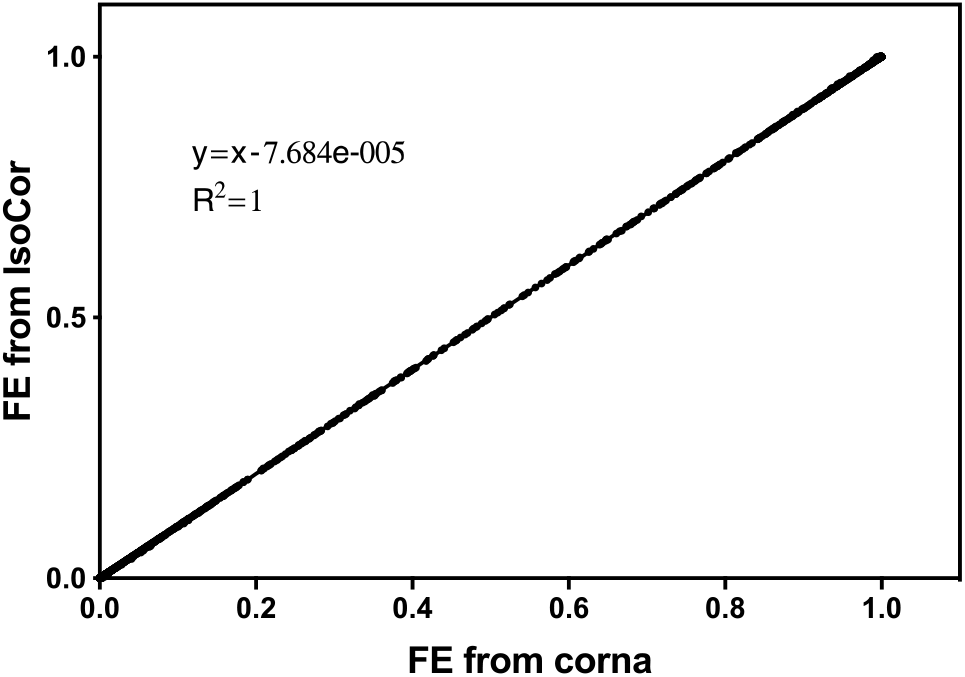
Comparison of natural abundance corrected intensities between IsoCorV1 and Corna with a large C13 labeled dataset

We also compared the results obtained after correcting a C13 labeled dataset with 24 samples containing 56 metabolites each, resulting in 7776 measurements. The output from these data were generated by processing the raw data [21] using El Maven 0.2.4 [22]. The peaks from El Maven were exported as group summary matrix format and the column names compound, label and formula were changed to Name, Label and Formula while keeping the sample columns as is before uploading to Corna. For IsoCorV1, the output was converted to batch processing .txt format using the example from the package as reference. These raw files were processed by both packages and output fractional enrichments were plotted with the output from IsoCorV1 on y-axis and from Corna on x-axis. From Figure 3, a slope of 1 between the fractional enrichment of metabolites calculated from natural abundance-corrected intensities from IsoCorV1 and Corna, can be observed with a shift of the order 10^−5^ indicating the outputs given by both packages are identical upto 5 decimal places.

Natural abundance correction of ultra high resolution MS data with both C13 and N15 tracers was compared with PyNAC which implements an iterative algorithm for dual isotope correction [9]. The example dataset from the PyNAC code repository, “13C15N_250DataSets.csv”, containing 250 samples with 18 metabolites each was used. The comparison of natural abundance corrected intensities between Corna and PyNAC is shown in Figure 4 (a). From the slope of the line fitted through the output values, it can be observed that the corrected values from PyNAC are slightly less than Corna which is expected because Corna also corrects for isotopes of resolved elements that were not measured. For example, if there is a hypothetical compound with the formula CHN, apart from correcting for natural abundance of C13 and N15, the label which was lost to natural abundance of H2 would also be corrected. This is done because even if the mass spectrometer is able to resolve H2 from C13 and N15, it is not measured by the design of the experiment, and therefore should be corrected (Supplementary S1.1.3). While the difference due to this correction is miniscule for hydrogen due to its small natural abundance, the difference will be significant when the element not measured in the experiment, such as C, has a significant natural abundance as shown in Figure 4 (b) and (c). The above scenario will occur if the experimental tracer is N15 or H2 and C13 is resolved from the tracer but not measured. AccuCor [17] implements single label correction by taking the above loss into account. Figure 4 (b) compares corrected intensities for deuterium labeled data for fructose-6-phosphate in sample 1_1 from the file “D_Sample_Input_Simple.xlsx” in AccuCor repository. While AccuCor and Corna return identical corrected values, PyNAC underestimates the corrected intensities. As AccuCor does not correct data with dual-tracers, results for dual label were compared by simulating data with 40% unlabeled uridine containing one N15 and one H2 isotope (Uridine-N15H2-label-1-1). The observed fractions were calculated by randomly choosing isotopes of the elements in uridine according to their natural abundance probabilities 10,000 times. The observed intensities were then corrected with Corna and PyNAC. As shown in Figure 4 (c), the expected experimental fractions are retrieved by Corna.

**Figure 4.**
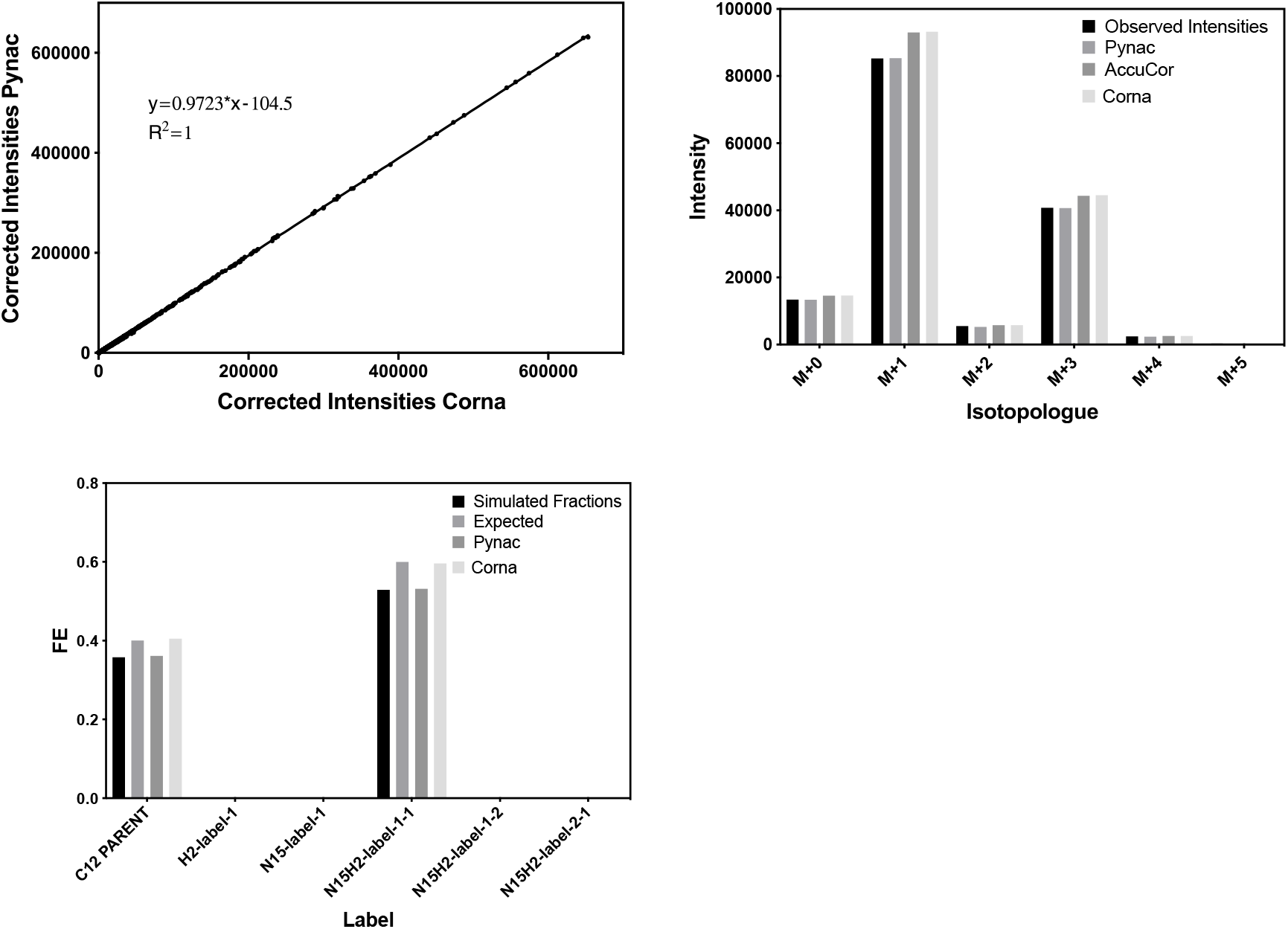
(a) Comparison of corrected values by PyNAC with those from Corna for an ultra high resolution dataset containing both C13 and N15 labels simultaneously (b) H2 labeled data for F6P corrected by PyNAC, AccuCor and Corna (c) H2 and N15 dual-labeled data simulated for Uridine with original expected fractions and corrected results from PyNAC and Corna

In MS measurements with derivatization, the atoms of the derivatizing agent contribute to the measured intensity of isotopologues and have to be accounted during natural abundance correction calculations [23]. GCMS data were validated with the algorithm implemented by Joanne K Kelleher et.al. (JKK matrix)^1^. A comparison of corrections for glucose derivatized as pentaacetate [24] is shown in Figure 5. The corrected intensities from JKK matrix agree with the corrected values from Corna. Another example with derivatizing agent TMS (Si(CH_3_)_4_) was validated with a theoretical distribution and is available in the Ipython notebook N4.

**Figure 5.**
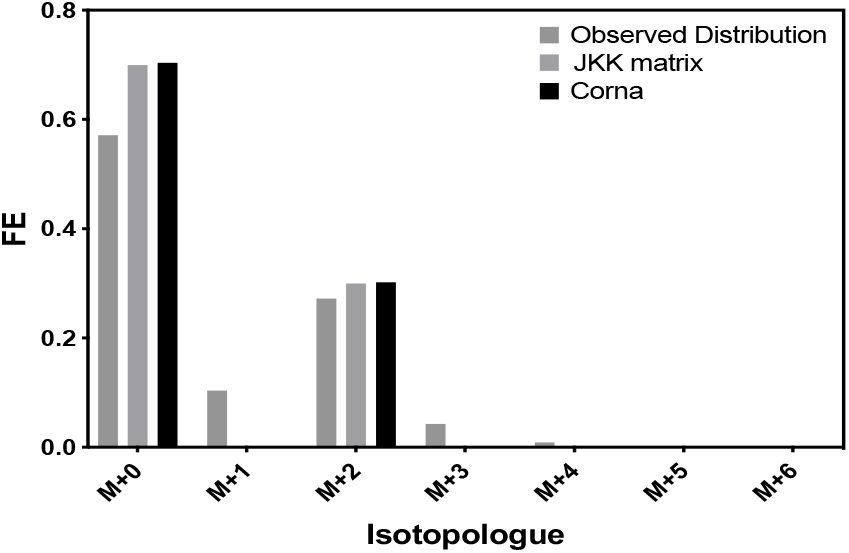
Natural abundance correction of C13 labeled glucose derivatized as pentaacetate using JKK matrix and Corna

Natural abundance correction for C13 labeled MS/MS data was done by the method described by Alves et. al.[20]. It takes into account the multiple labeling states of the daughter fragment given a particular label in the parent molecule. The data from a time course experiment with 7 time points containing 6 replicates each of INS-1 cells incubated in 4 different uniformly labeled C13 glucose concentrations were taken. The data were validated for the fragments of phosphoenolpyruvate (PEP) with mass 79 (PEP 167/79), pyruvate with mass 87 (Pyruvate 87/87), malate with mass 115 (Malate 133/115), aspartate with mass 88 (Aspartate 132/88), glutamate with mass 41 (Glutamate 146/41), citrate with mass 67 (Citrate 191/67), and succinate with mass 73 (Succinate 117/73). In addition to natural abundance correction, the authors also performed an additional correction, termed as background correction, using the intensities of the samples at 0 minutes, i.e., just before the labeled tracer element was introduced. The excel results of natural abundance correction and background correction were compared with the results from Corna and were found to be in complete agreement. Figure 6 shows the comparison for average corrected intensities of malate in INS-1 cells for six replicate samples at 240 minutes treated with [U-13C6] glucose at a concentration of 9 mM.

**Figure 6.**
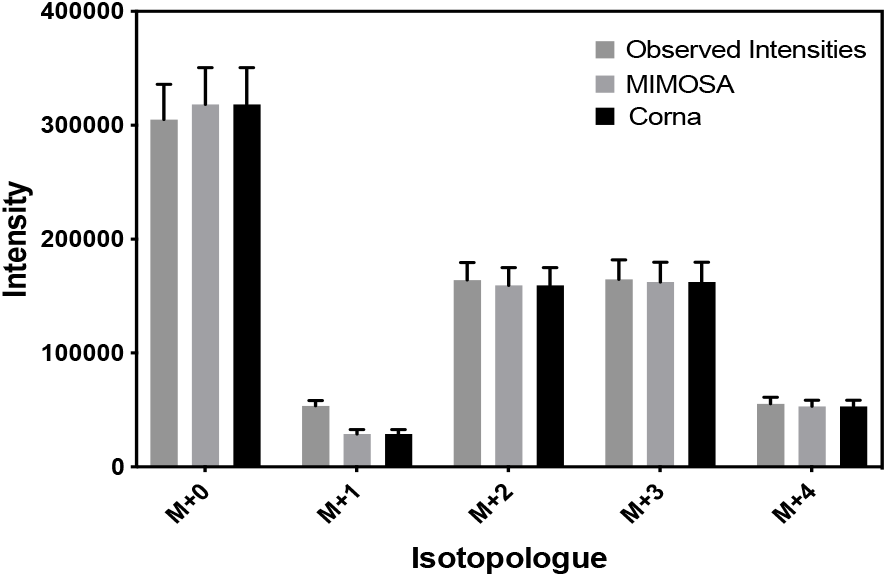
Comparison of C13 labeled Malate 133/115 fragment corrected with Corna and Microsoft Excel sheet implementation of correction algorithm from Alves Tiago et al.

A unique feature implemented in Corna is the auto-detection of indistinguishable isotopes based on the resolution of the instrument. The auto-detection algorithm was validated using the results from AccuCor [17] that can detect indistinguishable isotopes for Orbitrap given C13, N15 or H2 tracer elements. Intensity data for glutathione disulfide for sample N15_50_140k_A from the file “N_Sample_Input_Simple.xlsx” in the AccuCor package were corrected for natural abundance assuming three different resolutions defined at mass 200 for Orbitrap with 100% purity of tracer. In Figure 7 (a), it can be observed that the results for Corna and AccuCor are in complete agreement at all resolutions. As expected, different instrument resolutions give different corrected intensities depending on the isotopes resolved from the tracer element. When there are dual tracers, Corna detects indistinguishable isotopes separately for each tracer and generates a corresponding correction matrix. For example, an Orbitrap with minimal nominal resolution of 24500 at mass 200 can resolve H2 from N15 in purine with formula C5H4N4 but can not resolve C13 from H2. Corna can accommodate this scenario by treating C13 as indistinguishable from H2 but completely distinguishable from N15. To validate the correction for dual-tracers, simulated data were generated for 30% unlabeled purine with one H2 and one N15 isotope labeled experimentally at a resolution of 24500 defined at an m/z of 200. Figure 7 (b) shows that the 70% experimental label is recovered after correction in Corna.

**Figure 7.**
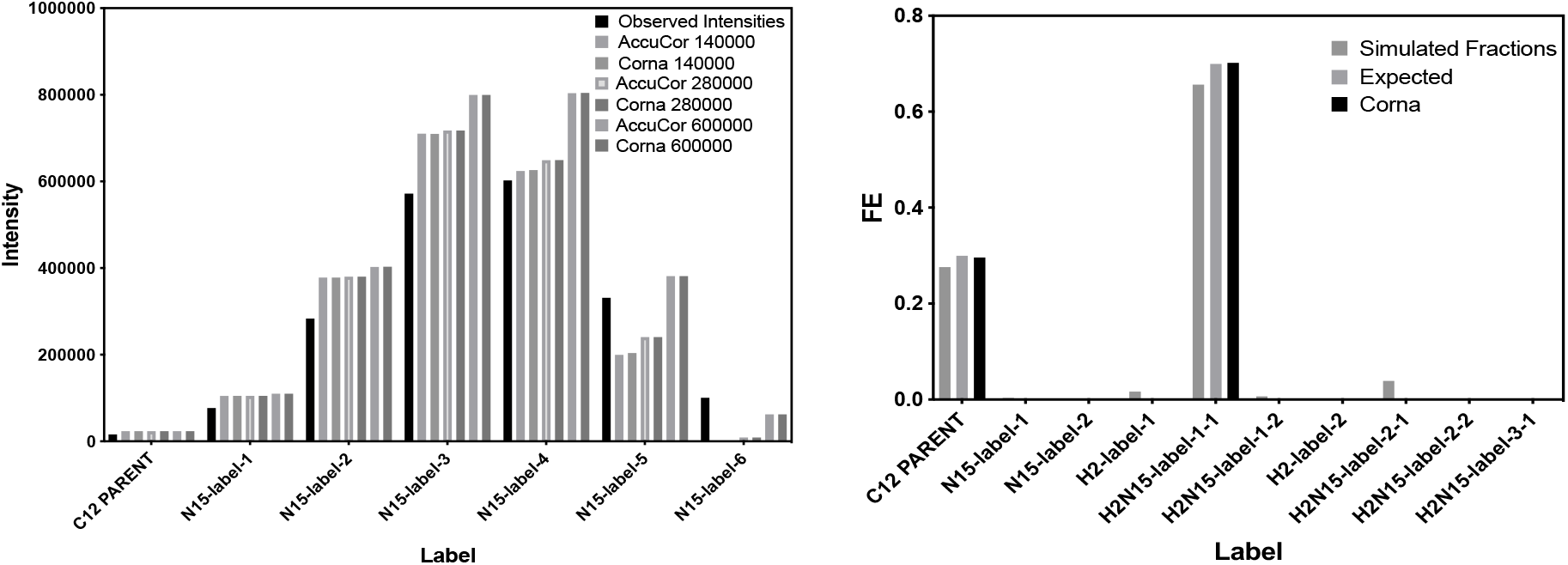
(a) Natural abundance correction of N15 labeled Glutathione Sulfide at 140000, 280000 and 600000 resolutions defined at m/z of 200 by AccuCor and Corna (b) Natural abundance correction of simulated purine values dual-labeled by H2 and N15 by Corna retrieves expected values

### Case Study: Integration of Corna with an open-source software, El-MAVEN

We present a case study to demonstrate the feasibility of using El-MAVEN, an open-source tool for integrating raw mass spectrometry data, and Corna together to analyze the raw LC-MS/MS data from Alves et. al. 2015. The results are validated against the original data which were processed wirh MultiQuant™ (Sciex) and Microsoft Excel (2013). Mass isotopologue data for several glycolytic and TCA cycle metabolites were measured over a time course of 7 time points with 6 replicates each i.e. a total of 42 samples. Representative metabolites with high signal to noise ratio and a negligible baseline (Malate and Glutamate) and a prominent baseline with low signal to noise ratio (Pyruvate and Lactate) were selected, amounting to 798 peaks. The total processing time with El-MAVEN followed by Corna was ~30 minutes as opposed to ~5.5 hours with MultiQuant™ and Microsoft Excel. Enrichment calculation took ~3 hours using Microsoft Excel but 6-fold less (~30 minutes) using Corna including the time required for creating sample metadata files, which is a one-time process. The total processing time will further reduce to 2-3 minutes if the metadata file creation is automated. The processing time increases in Microsoft Excel with increasing number of metabolites, but will not change in Corna. The averages and standard deviations of isotopologue fractions calculated from the first workflow are comparable to the published results as shown in Figure 8. The El-MAVEN and Corna pipeline is also vendor-neutral, open source, and available on GitHub, which makes it accessible and flexible for the user. Overall, this pipeline provides faster, accurate, and reproducible results that are less prone to manual errors.

**Figure 8.**
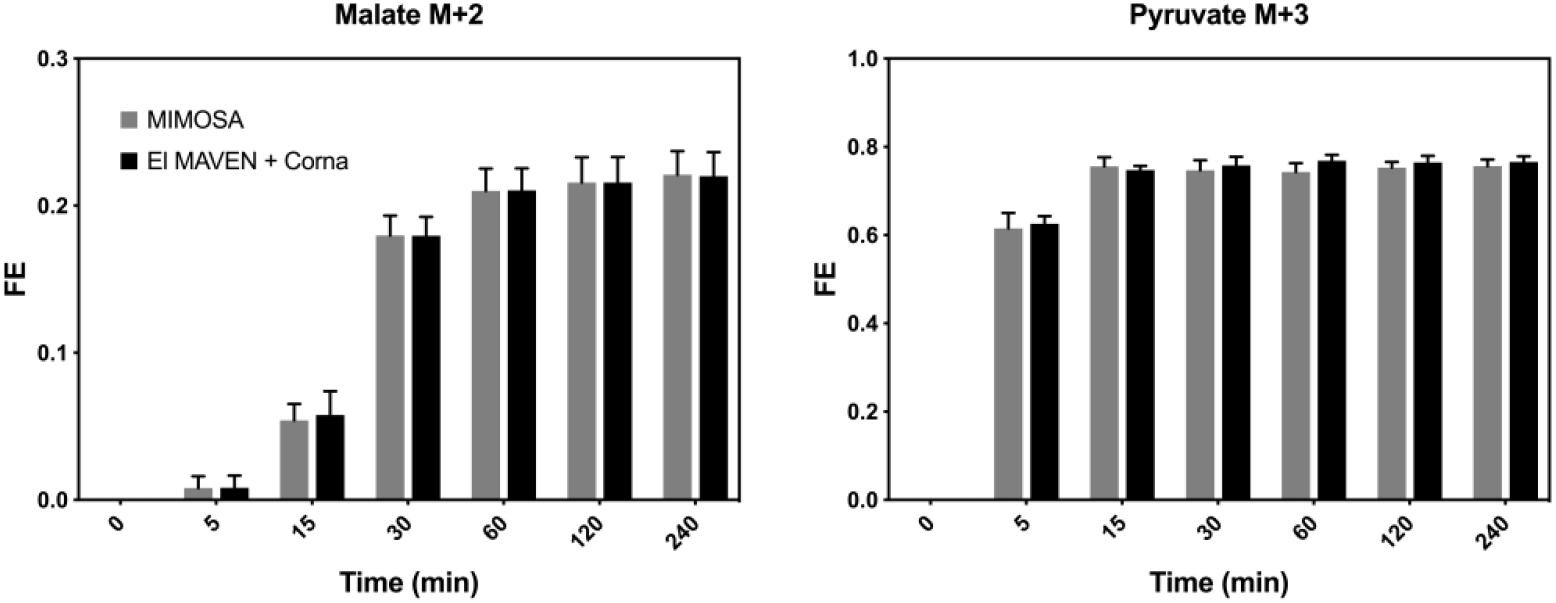
Comparison of fractional enrichments of Pyruvate M+3 and Malate M+2 using integrated El-MAVEN and Corna workflow with results from Alves et al. (2015)

## Discussion and Conclusion

Due to the increasing use of stable isotope tracers in studying metabolism, natural abundance correction has become a crucial step in data processing. The natural abundance correction algorithm used in a given stable isotopic tracer experiment, however, will have subtle differences depending on factors such as type of experimental technique, i.e., LCMS, LCMS-MS and GCMS, the resolution of the instrument, the type of stable isotopic tracer element, and the number of tracer elements. Typically, different algorithms for these experimental factors exist as standalone packages written in a variety of programming languages with their custom input and output formats that differ significantly. While designing a stable isotope tracer experiment, depending on the research question and availability of resources, the same lab might be involved in analyzing data from many different experimental techniques and methods. Since each tool has its own learning curve and input format, it can become very tedious to use different tools to correct natural abundance for different experiments. With Corna we have tried to address some of the above limitations. We have focused on combining different natural abundance correction algorithms in one place with similar architecture to make the processing easy and similar for different kinds of data. We hope that this integration will let the user focus on the data instead of worrying about learning a new tool or programming language. Using the same tool for different datasets also brings consistency and reproducibility in research.

Corna is written in Python, which is free and open source. Advanced users are welcome to contribute to the code. We provide commonly used workflows as Ipython notebooks for researchers with non-computational background. In Corna, input formats are separated from algorithms and the format is compatible with open-source peak integration software El-MAVEN [22]. Additional input formats can easily be supported in the future. Each sample in an experiment has important metadata which is lost while processing in every available tool that we know for natural abundance correction. With our focus on the user, we track sample metadata provided by the user and collate it with the final results, which reduces the time spent on combining sample information before and after natural abundance correction.

A new feature in Corna is the ability to auto-detect indistinguishable isotopes in experiments with dual tracers. Some high resolution mass spectrometer instruments are able to distinguish isotopes like C13 from N15. The packages available for performing natural abundance correction on these data assume infinite resolution, which means that it is assumed that if C13 can be resolved from N15 at all masses, it can also be resolved from other isotopes like H2. However, this assumption becomes invalid as the resolution of the machine varies with the mass of the metabolite.[17]. Corna does not assume ultra-high resolution and corrects for partial indistinguishability of isotopic elements in the data.

While Corna addresses some of the existing limitations with natural abundance correction tools and algorithms, there are features that remain to be incorporated. Isotope labels from vendors often come with a predefined purity of less than 100%. Corna currently does not take the purity of tracer into account. For MS-MS data, Corna is unable to handle low resolution measurements and only deals with single isotope tracers. In high resolution MS data, it is possible to resolve an isotopologue from another even when individual elements are not resolved separately [14], but this case is not yet handled in Corna. We hope to expand the available algorithms in Corna in the future to address the above limitations.

Algorithms and software packages to correct natural abundance have been around for a long time. But in the current era of technology when the bottleneck has shifted from generating data to processing it, it is crucial to recognize the importance of user experience and integrated and reproducible workflows. Corna is designed to work for the most commonly used isotope tracer experiments and can be plugged into custom workflows as per users requirements.

## Acknowledgements

This work was supported by NIH/NIDDK R01 DK-110181 (NCE) and DK-108283 (NCE).

## Footnotes

1 Course Material provided by Joanne K Kelleher during 10th Annual Course on Isotope Tracers in Metabolic Research: Principles and Practice of Kinetic Analysis, 2017

## References

[1] A. Chokkathukalam, D. Kim, M. P. Barrett, and D. J. Creek, “Europe PMC Funders Group Stable isotope-labeling studies in metabolomics : new insights into structure and dynamics of metabolic networks,” Bioanalysis, vol. 6, no. 4, pp. 511–524, 2014.

[2] F. S. Midani, M. L. Wynn, and S. Schnell, “The importance of accurately correcting for the natural abundance of stable isotopes,” Analytical Biochemistry. 2017.

[3] J. I. Brauman, “Least Squares Analysis and Simplification of Multi-Isotope Mass Spectra,” Anal. Chem., vol. 38, no. 4, pp. 607–610, 1966.

[4] J. Blackburn, “Computer Program for Multicomponent Spectrum Analysis Using Least Squares Method.,” Anal. Chem., vol. 37, no. 8, pp. 1001–1003, 1965.

[5] W.-N. P. Lee, L. O Byerley, E. A. Bergner, and J. Edmond, “Mass Isotopomer Analysis : Theoretical and Practical Considerations,” Biol. Mass Spectrom., vol. 20, 1991.

[6] C. A. Feroandez, C. Des Rosiers, S. F. Previs, F. David, and H. Brunengrabert, “Correction of ’ 3C Mass Isotopomer Distributions for Natural Stable Isotope Abundance,” J. MASS Spectrom., vol. 31, pp. 255–262, 1996.

[7] W. A. Van Winden, C. Wittmann, E. Heinzle, and J. J. Heijnen, “Correcting mass isotopomer distributions for naturally occurring isotopes,” Biotechnol. Bioeng., vol. 80, no. 4, pp. 477–479, 2002.

[8] S. A. Wahl, M. Dauner, and W. Wiechert, “New Tools for Mass Isotopomer Data Evaluation in 13C Flux Analysis: Mass Isotope Correction, Data Consistency Checking, and Precursor Relationships,” Biotechnol. Bioeng., 2004.

[9] H. N. B. Moseley, “Correcting for the effects of natural abundance in stable isotope resolved metabolomics experiments involving ultra-high resolution mass spectrometry,” BMC Bioinformatics, vol. 11, 2010.

[10] A. Rantanen, J. Rousu, J. T. Kokkonen, V. Tarkiainen, and R. A. Ketola, “Computing positional isotopomer distributions from tandem mass spectrometric data,” Metab. Eng., vol. 4, no. 4, pp. 285–294, 2002.

[11] S. Niedenführ, A. ten Pierick, P. T. N. van Dam, C. A. Suarez-Mendez, K. Nöh, and S. A. Wahl, “Natural isotope correction of MS/MS measurements for metabolomics and13C fluxomics,” Biotechnol. Bioeng., vol. 113, no. 5, pp. 1137–1147, 2016.

[12] C. Jungreuthmayer, S. Neubauer, T. Mairinger, J. Zanghellini, and S. Hann, “ICT: Isotope correction toolbox,” Bioinformatics, vol. 32, no. 1, pp. 154–156, 2015.

[13] P. Millard, F. Letisse, S. Sokol, and J. C. Portais, “IsoCor: Correcting MS data in isotope labeling experiments,” Bioinformatics, vol. 28, no. 9, pp. 1294–1296, 2012.

[14] P. Millard, B. Delépine, M. Guionnet, M. Heuillet, F. Bellvert, and F. Létisse, “IsoCor: isotope correction for high-resolution MS labeling experiments,” Bioinformatics, 2019.

[15] S. Trefely, P. Ashwell, and N. W. Snyder, “FluxFix: Automatic isotopologue normalization for metabolic tracer analysis,” BMC Bioinformatics, vol. 17, no. 1, pp. 1–8, 2016.

[16] W. Carreer, R. Flight, and H. Moseley, “A Computational Framework for High-Throughput Isotopic Natural Abundance Correction of Omics-Level Ultra-High Resolution FT-MS Datasets,” Metabolites, vol. 3, no. 4, pp. 853–866, 2013.

[17] X. Su, W. Lu, and J. D. Rabinowitz, “Metabolite Spectral Accuracy on Orbitraps,” Anal. Chem., vol. 89, no. 11, pp. 5940–5948, 2017.

[18] A. Jha and S. Pathak, “Polly,” 2017.

[19] A. G. Marshall, C. L. Hendrickson, and G. S. Jackson, “Fourier transform ion cyclotron resonance mass spectrometry: A primer,” Mass Spectrom. Rev., vol. 17, no. 1, pp. 1–35, 1998.

[20] T. C. Alves et al., “Integrated, Step-Wise, Mass-Isotopomeric Flux Analysis of the TCA Cycle,” Cell Metab., 2015.

[21] E. Melamud, L. Vastag, and J. D. Rabinowitz, “Metabolomic analysis and visualization engine for LC – MS data,” Anal. Chem., vol. 82, no. 23, pp. 9818–9826, 2010.

[22] S. P. Shubhra Agrawal, Sahil Kumar, Raghav Sehgal, Sabu George, Rishabh Gupta, Surbhi Poddar, Abhishek Jha, “El-MAVEN: A Fast, Robust, and User-Friendly Mass Spectrometry Data Processing Engine for Metabolomics,” in High-Throughput Metabolomics, vol. 1978, 2019, pp. 301–321.

[23] J. Rosenblatt, D. Chinkes, M. Wolfe, and R. R. Wolfe, “Stable isotope tracer analysis by GC-MS, including quantification of isotopomer effects.,” Am. J. Physiol., vol. 263, no. 3 Pt 1, pp. E584–96, 1992.

[24] G. W. Cline and G. I. Shulman, “Mass and positional isotopomer analysis of glucose metabolism in periportal and pericentral hepatocytes,” J. Biol. Chem., vol. 270, no. 47, pp. 28062–28067, 1995.

