## Supplementary for ">Corna - An Open Source Python Tool For Natural Abundance Correction In Isotope Tracer Experiments"

**S1. Algorithms**

Natural abundance correction for various mass spectrometry methods and resolutions have been described and implemented separately in the literature. These algorithms derive from the same basic concepts and here we try to describe the algorithms with those concepts.

**S1.1 MS**

**S1.1.1 Compound with Single tracer**

Consider a C_2_ molecule labeled by C13. The natural abundance of C13 is given by p and C12 by 1-p. Let M0 be the mass of the compound without any label, M1 with one increase, M2 with two and so on. M_obs_ is the intensity that is measured by the mass spectrometer corresponding to mass M and can contain both label from the experiment as well as naturally abundant isotope, M_corr_ is the expected intensity that contains label only from the experiment. M_obs_ can be written as function of M_corr_ as follows:

M0_obs_ = Both carbons must be C12 = Choosing 0 naturally abundant C13 atoms from 2 carbon atoms = $\binom{2}{0}p^{0}{(1-p)}^{2-0}$ M0_corr_ = (1-p)*(1-p)*M0_corr_

M1_obs_ = One C12 and One Natural C13 or One Experimental C13 and the other C12 = $\binom{2}{1}p^{1}{(1-p)}^{2-1}{M0}_{corr}+ \binom{2-1}{1-1}p^{0}{(1-p)}^{1-0}{M1}_{corr}=$2*(1-p)*p* M0_corr_ + (1-p)* M1_corr_

Here, in M1_corr_ we have no space left for natural abundant C13 after filling it with experimental C13 because total label in the system is 1.

M2_obs_ = Both Natural C13 or One Experimental C13 One natural C13 or Both Experimental C13 = $\binom{2}{2}p^{2}{(1-p)}^{2-2}{M0}_{corr}+ \binom{2-1}{2-1}p^{1}{(1-p)}^{1-1}{M1}_{corr}+ \binom{2-2}{2-2}p^{0}{(1-p)}^{0}{M2}_{corr}=$ p*p* M0_corr_ + p* M1_corr_ + 1* M2_corr_

The above problem is similar to choosing k C13 naturally abundant atoms with n being the total numberof C
atoms. k varies with number of already labeled C atoms from the experiment and total number of observed labeled atoms.

**S1.1.2 Compound with Single tracer and indistinguishable isotopes (low resolution)**

Now, consider a hypothetical molecule CH where H2 is indistinguishable with C13 i.e M+1 can be mass added due to H2 (deuterium) or C13. If natural abundance of H2 is h and we consider only two isotopes of Hydrogen are possible than abundance of H1 would be (1-h)

M0_obs_ = Both C12 and H1 = Choose 0 naturally abundant C13 from one Carbon and 0 naturally abundant D2 from one Hydrogen = $\binom{1}{0}p^{0}{(1-p)}^{1-0}$ $\binom{1}{0}h^{0}{(1-h)}^{1-0}$ M0_corr_= (1-p)*(1-h)* M0_corr_

M1_obs_ = One natural C13 and One natural H1or One natural H2 and one natural C12 or One Experimental C13 and one natural H1 = $(\binom{1-0}{0-0}p^{0}{(1-p)}^{1-0}$ $\binom{1-0}{1-0}h^{1}{(1-h)}^{1-1}$ + $\binom{1-0}{1-0}p^{1}{(1-p)}^{1-1}$ $\binom{1-0}{0-0}h^{0}{(1-h)}^{1-0})*$ M0_corr_ + $\binom{1-1}{1-1}p^{0}{(1-p)}^{0-0}$ $\binom{1-0}{0-0}h^{0}{(1-h)}^{1-0}*{M1}_{corr}=$ (p*(1-h) + h*(1-p))* M0_corr +_ (1-h)*M1_corr_

The above problem is also similar to choosing k items from n, where H2 can only be chosen from number of H available and C13 from number of C available. As we are not able to distinguish if the increase in mass is due to H2 or C13, we have to consider both cases where it is coming from H2 or C13.

**S1.1.3 Compound with Single tracer and resolved isotopes (ultra high resolution)**

In the same hypothetical molecule CH if H2 was resolved from C13 i.e. the mass spectrometer can tell whether it is M+1 from C13 or M+1 from H2, while measuring we only integrate the peak due to C13. Now the equations would change slightly from what we had in the previous section:

M0_obs_ = Both C12 and H1 = Choose 0 naturally abundant C13 from one Carbon and 0 naturally abundant D2 from one Hydrogen = $\binom{1}{0}p^{0}{(1-p)}^{1-0}$ $\binom{1}{0}h^{0}{(1-h)}^{1-0}$ M0_corr_= (1-p)*(1-h)* M0_corr_

M1_obs_ = One natural C13 and One natural H1 or One Experimental C13 and one natural H1 = $(\binom{1-0}{0-0}p^{0}{(1-p)}^{1-0}$ $\binom{1-0}{1-0}h^{1}{(1-h)}^{1-1})*$ M0_corr_ + $\binom{1-1}{1-1}p^{0}{(1-p)}^{0-0}$ $\binom{1-0}{0-0}h^{0}{(1-h)}^{1-0}*{M1}_{corr}=$ p*(1-h) * M0_corr +_ (1-h)*M1_corr_

The system still has to be corrected for the natural abundance of resolved isotopes but now we don’t have to worry about anything being measured with the desired tracer.

**S1.1.4 Compound with Single tracer and not experimentally labeled tracer atoms (derivatizer)**

In some experiments, the compound is derivatized by combining with other compound to increase the volatility of the substance to be measured. As the adduct is not part of the labeling experiment its atoms can’t be labeled. For example, let us say we have a compound with molecular formula C. It is experimentally labeled so we can have label going up to M1, and now before measuring we derivatize it to C_2_, so there is an additional C atom which would bring its natural abundance but it can’t be labeled from the experiment. In this scenario the equations will be:

M0_obs_ = Both carbons must be C12 = Choosing 0 naturally abundant C13 atoms from 2 carbon atoms = $\binom{2}{0}p^{0}{(1-p)}^{2-0}$ M0_corr_ = (1-p)*(1-p)*M0_corr_

M1_obs_ = One C12 and One Natural C13 or One Experimental C13 and the other C12 = $\binom{2}{1}p^{1}{(1-p)}^{2-1}{M0}_{corr}+ \binom{2-1}{1-1}p^{0}{(1-p)}^{1-0}{M1}_{corr}=$2*(1-p)*p* M0_corr_ + (1-p)* M1_corr_

Now because the label can only be upto M1, we would stop here. It should be noticed that these are the same first two equations from the section “Compound with Single tracer“. So we can correct for such compounds by assuming all atoms of tracer can be labeled (2 in this case) but restricting the equations to maximum label possible according the original compound (which is 1 here).

All the system of equations above can be solved iteratively for M_corr_ values given M_obs_ or the system can be represented with a correction matrix (CM):

M_obs_ = CM * M_corr_

Which can be solved to give M_corr_ by:

M_corr_ = CM^-1^ * M_obs_

**S1.1.5 Compound with Dual tracer and resolved isotopes (ultra high res)**

The same principles can be extended for dual tracers as shown here with a hypothetical compound CHN labeled experimentally by C13 and N15. Now depending on the label instead of M+1 we would have M+1 with C13 (M_C13_1_obs_) or M+1 with N15 (M_N15_1_obs_) and so on. Here we write down the equations for two tracers assuming the natural abundance to be c,n and h respectively for C13, N15 and H2:

M0_obs_ = (1-c)(1-n)(1-h) * M0_corr_

M_C13_1_obs_ = (1-n)(1-h) * M_C13_1_corr_ + c(1-n)(1-h) * M0_corr_

M_N15_1_obs_ = (1-c)(1-h) * M_N15_1_corr_ + n(1-c)(1-h) * M0_corr_

M_C13_N15_1_1_obs_ = (1-h) * M_C13_N15_1_1_corr_ + n(1-h) * M_C13_1_corr_ + c(1-h) * M_N15_1_corr_ + nc(1-h) * M0_corr_

The matrix equivalent of this is sequentially multiplying with each label matrix and then the resolved isotope:

$\frac{\left[ \begin{matrix} (1-c)M0corr \\ cM0corr + M\_C13\_1corr \end{matrix} \right] = \left[ \begin{matrix} 1-c & 0 \\ c & 1 \end{matrix} \right] * \left[ \begin{matrix} M0corr \\ M\_C13\_1corr \end{matrix} \right]}{\left[ \begin{matrix} (1-c)M\_N15\_1corr \\ cM\_N15\_corr + M\_C13\_N15\_1\_1corr \end{matrix} \right] \left[ \begin{matrix} 1-c & 0 \\ c & 1 \end{matrix} \right] * \left[ \begin{matrix} M\_N15\_1corr \\ M\_C13\_N15\_1\_1corr \end{matrix} \right]}$

$\frac{\left[ \begin{matrix} \left( 1-n \right)\left( 1-c \right)M0corr \\ n\left( 1-c \right)M0corr+\left( 1-c \right)M_{N{15}_{1corr}} \end{matrix} \right] = \left[ \begin{matrix} 1-n & 0 \\ n & 1 \end{matrix} \right] * \left[ \begin{matrix} (1-c)M0corr \\ (1-c)M\_N15\_1corr \end{matrix} \right]}{\left[ \begin{matrix} c(1-n)M0corr + (1-n)M\_C13\_1corr \\ M\_C13\_N15\_1\_1corr + nM\_C13\_1corr + cM\_N15\_1corr + ncM0corr \end{matrix} \right]\left[ \begin{matrix} 1-n & 0 \\ n & 1 \end{matrix} \right] * \left[ \begin{matrix} cM0corr + M\_C13\_1corr \\ cM\_N15\_corr + M\_C13\_N15\_1\_1corr \end{matrix} \right]}$

$\frac{\left[ \begin{matrix} M0obs \\ M\_N15\_1\mathrm{obs} \end{matrix} \right] = \left[ \begin{matrix} 1-h & 0 \\ 0 & 1-h \end{matrix} \right] * \left[ \begin{matrix} (1-n)(1-c)M0corr \\ n(1-c)M0corr+(1-c)M\_N15\_1corr \end{matrix} \right]}{\left[ \begin{matrix} M\_C13\_1\mathrm{obs} \\ M\_C13\_N15\_1\_1obs \end{matrix} \right] \left[ \begin{matrix} 1-h & 0 \\ 0 & 1-h \end{matrix} \right] * \left[ \begin{matrix} c(1-n)M0corr + (1-n)M\_C13\_1corr \\ M\_C13\_N15\_1\_1corr + nM\_C13\_1corr + cM\_N15\_1corr + ncM0corr \end{matrix} \right]}$

**S1.1.6 When constructing a matrix is not trivial**

Compound with Single tracer and partially resolved isotopes (autodetect)

Considering a compound like CNH_2_ labeled with H2, it is possible that C13 and N15 are indistinguishable with H2

M0_obs_ = (1-c)(1-n)(1-h)^2^ * M0_corr_

M1_obs_ = ((1-n)(1-c)*2h(1-h) + c(1-n)(1-h)^2^ + n(1-c) (1-h)^2^ ) * M0_corr_ + (1-c)(1-h)(1-n) * M1_corr_

M2_obs_ = (h^2^(1-c)(1-n) + 2h(1-h)c(1-n) + 2h(1-h)n(1-c) + cn(1-h)^2^)* M0_corr_ + ((1-c)(1-n)h + c(1-n)(1-h) + (1-c)n(1-h)) * M1_corr_ + (1-c)(1-n)M2_corr_

The correction matrix for this system of equation is:

$\left[ \begin{matrix} (1-c)(1-n){(1-h)}^{2} & 0 & 0 \\ \left( c+n-2cn \right){(1-h)}^{2}+2h(1-h)(1-n)(1-c) & (1-c)(1-n)(1-h) & 0 \\ cn{(1-h)}^{2}+\left( c+n-2cn \right)2h\left( 1-h \right)+h^{2}(1-c)(1-n) & \left( c+n-2cn \right)\left( 1-h \right)+h(1-c)(1-n) & (1-c)(1-n) \end{matrix} \right]$

Simplified to:

$$\left[ \begin{matrix} (1-c)(1-n) & 0 & 0 \\ c+n-2cn & (1-c)(1-n) & 0 \\ cn & c+n-2cn & (1-c)(1-n) \end{matrix} \right] \times\left[ \begin{matrix} {(1-h)}^{2} & 0 & 0 \\ 2h(1-h) & 1-h & 0 \\ h^{2} & h & 1 \end{matrix} \right]$$

And then:

$$\left[ \begin{matrix} 1-c & 0 & 0 \\ c & 1-c & 0 \\ 0 & c & 1-c \end{matrix} \right]\times\left[ \begin{matrix} 1-n & 0 & 0 \\ n & 1-n & 0 \\ 0 & n & 1-n \end{matrix} \right]\times\left[ \begin{matrix} {(1-h)}^{2} & 0 & 0 \\ 2h(1-h) & 1-h & 0 \\ h^{2} & h & 1 \end{matrix} \right]$$

But, it is possible to have a scenario where when taken separately C13 and N15 are indistinguishable from H2, taken together they are not i.e. i.e. C13N15 can be resolved from H2_2_ (IsoCor V2) which would mean our initial matrix becomes:

$$\left[ \begin{matrix} (1-c)(1-n){(1-h)}^{2} & 0 & 0 \\ \left( c+n-2cn \right){(1-h)}^{2}+2h(1-h)(1-n)(1-c) & (1-c)(1-n)(1-h) & 0 \\ cn{(1-h)}^{2}+\left( c+n-2cn \right)2h\left( 1-h \right)+h^{2}(1-c)(1-n) & \left( c+n-2cn \right)\left( 1-h \right)+h(1-c)(1-n) & (1-c)(1-n) \end{matrix} \right]$$

Which can be simplied to:

$$\left[ \begin{matrix} (1-c)(1-n) & 0 & 0 \\ c+n-2cn & (1-c)(1-n) & 0 \\ cn & c+n-2cn & (1-c)(1-n) \end{matrix} \right] \times\left[ \begin{matrix} {(1-h)}^{2} & 0 & 0 \\ 2h(1-h) & 1-h & 0 \\ h^{2} & h & 1 \end{matrix} \right]$$

Further simplication of this matrix is not trivial, hence for dealing with such cases it might be better to get each term of the matrix individually or solve the equations iteratively.

Compound with Dual tracer and isotopes indistinguishable with both

In the scenario where while C13 and N15 are resolved from each other but H2 can’t be distinguished from either, for the hypothetical compound CHN the equations will be:

M0_obs_ = (1-c)(1-n)(1-h)M0_corr_

M_C13_1_obs_ = (1-n)(1-h)M_C13_1_corr_ + c(1-n)(1-h)M0_corr_ + h(1-c)(1-n)M0_corr_

M_N15_1_obs_ = (1-c)(1-h)M_N15_1_corr_ + n(1-c)(1-h)M0_corr_ + h(1-c)(1-n)M0_corr_

M_C13_N15_1_1_obs_ = (1-h)M_C13_N15_1_1_corr_ + n(1-h) M_C13_1_corr_ + c(1-h) M_N15_1_corr_ + h(1-c) M_N15_1_corr_ + h(1-n)M_C13_1_corr_ + nc(1-h)M0_corr_ + nh(1-c)M0_corr_ + ch(1-n)M0_corr_

Decomposing these equations into matrices doesn’t seem to be trivial either and a better solution would be to generate each term and solve the system of equations.

Corna doesn’t handle the above two scenarios as of now.

**S1.2 MS/MS**

For MSMS, the principal of choosing natural abundant carbon atoms is the same, the only difference being that choices of natural abundant atoms are divided between the daughter fragments so we have to calculate probabilities for both and multiply to get a single coefficient as per the multiplication rule of probability. Here we explain the scenario with single tracer, C13.

If we have a fragment with D number of carbon atoms out of which l are experimentally labeled and the total label is j, the number of ways that we can choose positions of natural abundant C13 atoms is:

$$\binom{D-l}{j-l}$$

given p is the probability of naturally abundant C13, the probability of such an event is:

$$P\left( A \right)=\binom{D-l}{j-l}p^{j-l}\left( 1-p \right)^{D-j}$$

We have another fragment with N-D number of carbon atoms out of which k-l are experimentally labeled and the total label is i-j, the number of ways that we can choose positions of natural abundant C13 atoms is:

$$\binom{N-D-(k-l)}{i-j-(k-l)}$$

given p is the probability of naturally abundant C13, the probability of such an event is:

$$P\left( B \right)=\binom{N-D-(k-l)}{i-j-(k-l)}p^{i-j-(k-l)}\left( 1-p \right)^{N-D-(i-j)}$$

If we combine these fragments, we get a compound with N carbon atoms out of which k are experimentally labeled and i is the total label, and the probability of this happening is:

$$P\left( A\cap B \right)=P\left( A \right)\times P(B)$$

which is the probability of conversion of a fragment with parent label k and daughter label l, $M_{kl}$, to a fragment with parent label i and daughter label j, $M_{ij}$

Summing up such k,l where k and l are less than i and j, we get the following equation:

$$M_{ij}^{'}=\sum_{k,l<i,j} \binom{D-l}{j-l}\binom{N-D-(k-l)}{i-j-(k-l)}p^{i-k}{(1-p)}^{N-i}M_{kl}+{(1-p)}^{N-i}M_{ij}$$

$$M^{'} is the observed intensity$$

$$Mis the expected/corrected intensity$$

$$M_{ij} here i is the number of C13 atoms in parent,$$

$$j is the number of C13 atoms in daughter fragment$$

$$N is the total number of C atoms in the parent fragment$$

$$D is the total number of C atoms in the daughter fragment$$

$$p is the natural abundance of C13$$

The package implements a simpler version of this equation where higher order values of p are not considered because for C13 p = 0.011 which is <<1. Also, using p<<1 the above equation is solved to give the formula described in Tiago Alves et al:

$$M_{ij}=M_{ij}^{,}*\left( 1+p\left( N-i \right) \right)- M_{i-1j}^{'}*p\left( \left( N-D \right)-\left( i-j-1 \right) \right)- M_{i-1j-1}^{,}*p(D-\left( j-1 \right))$$

**S2. Extension of Tracers**

While the package was validated for C13, H2 and N15 tracers corna also supports other lesser used tracers like O17, O18, S33. As the most commonly used tracers in metabolomic studies are the former three, we focused on validating those. From the packages used for validation AccuCor and Pynac support only C13, N15 and H2. IsoCor has the option to specify other tracers like O17, O18, S33 but for tracers with +2 mass like O18 an error shows up when uploading data containing label more than n+1 where n is the number of atoms of the tracer. For +2 tracers one would expect the label to go till 2n. The correction matrix for O would be different if the label is O17 or O18 as explained below with an example of labeled H_2_O with O17 or O18 where H is fully resolved with O. In the following equations consider the natural abundance be o_16_, o_17_ and o_18_ respectively for O16, O17 and O18, and h_1_ and h_2_ respectively for H1 and H2.

O17 label

M0_obs_ = o_16_ (h_1_)^2^M0_corr_

M1_obs_ = (h_1_)^2^* M1_corr_ + o_17_ (h_1_)^2^M0_corr_

$$\begin{matrix} {M0}_{obs} \\ {M1}_{obs} \end{matrix} =\left[ \begin{matrix} h_{1}^{2} & 0 \\ 0 & h_{1}^{2} \end{matrix} \right] \times\left[ \begin{matrix} o_{16} & 0 \\ o_{17} & 1 \end{matrix} \right]\times\begin{matrix} {M0}_{corr} \\ {M1}_{corr} \end{matrix}$$

O18 label

M0_obs_ = o_16_ (h_1_)^2^M0_corr_

M1_obs_ = o_17_ (h_1_)^2^M0_corr_

M2 _obs_= (h_1_)^2^M2_corr_+ o_18_ (h_1_)^2^M0_cor_

$$\begin{matrix} \begin{matrix} {M0}_{obs} \\ {M1}_{obs} \end{matrix} \\ {M2}_{obs} \end{matrix} =\left[ \begin{aligned} \begin{matrix} h_{1}^{2} & 0 \\ 0 & h_{1}^{2} \end{matrix} \begin{matrix} 0 \\ 0 \end{matrix} \\ \begin{matrix} 0 & 0 & h_{1}^{2} \end{matrix} \end{aligned} \right] \times\left[ \begin{aligned} \begin{matrix} o_{16} & 0 \\ o_{17} & 0 \end{matrix} \begin{matrix} 0 \\ 0 \end{matrix} \\ \begin{matrix} o_{18} & 0 & 1 \end{matrix} \end{aligned} \right]\times\begin{matrix} \begin{matrix} {M0}_{corr} \\ {M1}_{corr} \end{matrix} \\ {M2}_{corr} \end{matrix}$$

Because M1_corr_ is 0, values in the matrix multiplied by this term (struck out above) can be ignored and in the final output are not relevant. Therefore, one way to create the matrix could be creating a 3x3 matrix distributing all the possible combinations of [o_16_, o_17,_ o_18_] irrespective of the label which would lead to some incorrect terms in the matrix but those would be multiplied by 0 and hence not affect the final output. The current algorithm in corna would output a matrix for O18 as follows:

$$\begin{matrix} \begin{matrix} {M0}_{obs} \\ {M1}_{obs} \end{matrix} \\ {M2}_{obs} \end{matrix} =\left[ \begin{aligned} \begin{matrix} h_{1}^{2} & 0 \\ 0 & h_{1}^{2} \end{matrix} \begin{matrix} 0 \\ 0 \end{matrix} \\ \begin{matrix} 0 & 0 & h_{1}^{2} \end{matrix} \end{aligned} \right] \times\left[ \begin{aligned} \begin{matrix} o_{16} & 0 \\ o_{17} & o_{16} \end{matrix} \begin{matrix} 0 \\ 0 \end{matrix} \\ \begin{matrix} o_{18} & o_{17} & 1 \end{matrix} \end{aligned} \right]\times\begin{matrix} \begin{matrix} {M0}_{corr} \\ {M1}_{corr} \end{matrix} \\ {M2}_{corr} \end{matrix}$$

which when inversed and multiplied should give correct final output as long as M1_obs_ is a zero input. But these results have to be validated yet.
